# CD34+ Stromal Cell/Telocytes Demonstrate a Dynamic Pattern of Distribution During Healing of Post-Infarcted Myocardium in Middle-Aged Sprague-Dawley Rats

**DOI:** 10.1101/2024.09.13.612962

**Authors:** Daniel T. Schneider, Eduard I. Dedkov

**Author notes:** **Corresponding Author:** Daniel T. Schneider.

## Abstract

**Introduction:** Myocardial CD34+ stromal cells/telocytes (SC/TCs) have been recently recognized as a novel resident cell which may play an important role in the repair process following acute myocardial infarction (MI). This study aims to determine the spatiotemporal dynamics of CD34+ SC/TCs within the left ventricular (LV) wall during the late inflammatory and proliferative phases of post-MI scar formation.

**Methods:** A large transmural MI was induced in middle-aged, Sprague-Dawley rats by permanent ligation of the left anterior descending coronary artery. To recognize proliferating cells, rats were infused with 5-bromo-2’-deoxyuridine (BrdU) in a dose of 12.5 mg/kg/day for 72 hours via intraperitoneal osmotic minipumps on day 0, 4, or 11 after surgery. The rats were euthanized on day 3, 7 and 14 after MI, and their hearts were processed for histology and immunostaining.

**Results:** Three days after MI, CD34+ SC/TCs were absent within the necrotic myocardial tissue but were visible around the surviving cardiac myocytes (CMs) bordering the infarcted region, including those remaining in subepicardial and subendocardial regions, and in the adventitia of residual coronary vessels. Seven days after MI, many of the CD34+ SC/TCs located at the periphery of the developing scar appeared enlarged and contained the BrdU labeling, indicating the cell proliferation. At the same time, elongated CD34+ SC/TCs, which lacked BrdU labeling, were noticed closer to the necrotic zone residing in the interstitial areas between the intact basement membranes left from resorbed CMs, suggesting their migratory activity. Fourteen days after MI, CD34+ SC/TCs were distributed throughout the entire post-infarcted region except for the areas occupied by necrotic tissue, myofibroblast-rich granulation tissue, and the fibroelastic thickenings of the endocardium affected by an MI. Furthermore, accumulated clusters of flattened CD34+ SC/TCs cells were apparent in the areas where the edges of surviving CMs extend into the fibrotic portion of the scar.

**Conclusion:** These findings, for the first time, demonstrate that a population of myocardial CD34+ SC/TCs follow a dynamic pattern of spatiotemporal distribution within the healing myocardium suggesting their direct involvement in post-MI repair process and scar formation.

## 1. INTRODUCTION

The sudden death of a substantial volume of left ventricular (LV) myocardium secondary to an acute myocardial infarction (MI) triggers a process involving LV wall scarring and chamber remodeling. Progressive, post-MI cardiac remodeling associated with scar thinning and expansion is implicated in the development of systolic heart failure (HF)^1,2^. Although the mechanisms behind the scar development and heart remodeling have been a target of extensive research for the last several decades, a comprehensive understanding of all involved contributors is far from complete.

Myocardial healing after an ischemic insult is well-defined and is comprised of both short- and long-term events that involve multiple cell lines and various chemical, cellular, and humoral mediators. Traditionally, the process is divided into three overlapping phases: (1) the inflammatory phase (approximately zero to three days post-MI), characterized by the immune reaction to damaged cells and the clearing of necrotic tissue by macrophages, (2) the reparative or proliferative phase (approximately three days to three weeks post-MI), characterized by the growth of granulation tissue rich with proliferating and collagen producing myofibroblasts, and (3) the maturation phase (>three weeks post-MI), characterized by the remodeling and compaction of the collagenous extracellular matrix that will serve as the basis for fibrotic scar tissue^3^.

Although the major players of myocardial wound healing process are long established, a fairly large number of modern-day studies have continued to hunt for the missing “piece of the puzzle” linking pathological changes in post-MI scar formation with the novel contributors that can be therapeutically targeted in the future. A recently identified population of cardiac CD34+ stromal cells/telocytes (SC/TCs) are considered one of these potentially vital targets for therapy^4^.

In several recent studies, the existence of CD34+ SC/TCs within the heart has been considered as a crucial element in myocardial tissue organization in development, as well as a fundamental supporter of myocardial tissue repair^5,6^. The mechanisms by which this type of cells could achieve such crucial roles have been thought to involve direct contributions to the angiogenic response, paracrine signaling of many resident and infiltrative cell types, and cardiac extracellular matrix remodeling^7,8^.

Some studies have also demonstrated that CD34+ SC/TCs are particularly sensitive to ischemia and are ubiquitously absent within the necrotic core of infarcted myocardium which eventually becomes a non-contractile, fibrotic scar^9^. Considering these observations, it has been proposed that the preservation or engraftment of CD34+ SC/TCs in ischemic myocardium could facilitate the formation of less fibrotic, better functioning tissue in the infarcted region. While the results from such experimental approach were initially promising, no definitive therapies have been successfully implemented to date^10^.

While there may be an incentive to leap immediately from simple abovementioned observations to clinical therapies, there remains a huge gap in understanding as to the ultimate role these cells may play within the heart. The problem this creates is that it limits the understanding of CD34+ SC/TCs to an effect modifier in tissue repair, when their role is likely a markedly complicated confounding variable given their known interactions with so many other cell types. As such, it is important to investigate CD34+ SC/TCs interactions with the individual components implicated in myocardial repair and regeneration, including the inflammatory response, development of granulation tissue, and subsequent fibrosis.

This study aims to characterize the spatiotemporal dynamics of CD34+ SC/TC’s within post-infarcted myocardium during formation of the post-MI scar, with a focus on the late inflammatory and proliferative phases of the healing process.

## 2. MATERIALS AND METHODS

### 2.1. Animals and Experimental Protocol

All animal procedures were approved by the Institutional Animal Care and Use Committee and performed in accordance with the *Guide for the Care and Use of Laboratory Animals, 8th edition*^11^.

A large transmural MI was induced in 12-month-old male (n = 24) Sprague-Dawley rats (Charles River Laboratories, Inc., Wilmington, MA) under ketamine (100 mg/kg intraperitoneal [i.p.]) and xylazine (10 mg/kg i.p.) anesthesia by permanent ligation of the left anterior descending (LAD) coronary artery, as previously detailed^12^. Following surgery, the rats were housed under climate-controlled conditions at a 12-hour light/dark cycle and provided with standard rat chow and water ad libitum.

On days 0, 4 and 11 after LAD artery ligation, three separate groups of rats (n = 6 to 7 per group) had received 5-bromo-2’-deoxyuridine (BrdU; Sigma, St. Louise, MO) in phosphate-buffered saline (PBS) via i.p. ALZET osmotic pumps (Durect, Cupertino, CA) at a dose 12.5 mg/kg/day for 72 hours, as previously detailed^13^. Subsequently, the rats in these groups were euthanized 3 days, 1 and 2 weeks after MI (n = 5 to 7 animals per group) and their hearts were collected for further evaluations. Briefly, rats were anesthetized with 4% isoflurane in pure oxygen and their hearts were arrested in diastole by an intracardiac injection with 100 mmol/L potassium chloride. Then the hearts were excised, perfusion-fixed on a Langendorff apparatus with 4% paraformaldehyde (PFA) in PBS for 20 minutes and were stored in a fresh solution of 4% PFA in PBS for 48 hours at +4°C, as previously detailed^14^. Subsequently, each heart was cut transversely into 2-mm-thick parallel slices using a blade guillotine and the two midventricular slices were further processed into paraffin blocks for histology and immunostaining.

### 2.2. Histology and Immunostaining

Transverse, 8.0-µm-thick serial sections were cut from all paraffin-embedded slices onto microscope slides. From each heart, several serial sections were routinely stained with hematoxylin and eosin (H&E) and Masson’s trichrome stains. Other serial sections were subjected to immunolabeling with various combinations of the primary antibodies (Table 1), secondary antibodies (Table 2), and lectins (Table 3) following ether enzymatic or heat-induced antigen retrieval methods. Deparaffinized and rehydrated sections were treated either with proteolytic enzymes, such as proteinase K (20 µg/mL; cat. no. P4850; Sigma, St. Louis, MO) and trypsin (0.025%; cat. no. SM-2001-C; Millipore Sigma, Burlington, MA) in PBS, for 15 minutes at +37°C or boiled in 0.01M sodium citrate buffer at pH 6.0 for 30 minutes followed by cooling for 20 minutes at room temperature.

**Table 1.** Primary Antibodies Used for Immunostaining.

| Table 1. Primary Antibodies Used for Immunostaining |  |  |  |  |  |  |  |
| --- | --- | --- | --- | --- | --- | --- | --- |
| Antigen Description | Host | Clone | Isotype | Antigen Retrieval | Dilution | Catalog No. | Source |
| Actin, a-smooth muscle | Mouse | 1A4 | IgG2a | PIER, HIER | 1:300 | A2547 | Sigma; St. Louis, MO |
| Bromodeoxyuridine | Mouse | 131-14871 | IgG1 | HIER | 1:1000 | MAB4072 | Millipore; Temecula, CA |
| CD34 | Rabbit | EP373Y | IgG | HIER | 1:100 | ab81289 | Abcam; Waltham, MA |

**Abbreviations:** a, alpha; Ag, antigen; DSHB, Developmental Studies Hybridoma Bank; HIER, heat-induced epitope (antigen) retrieval with citrate buffer (pH 6.0); PIER, protease-induced epitope (antigen) retrieval with proteinase K (20 µg/ml) or trypsin (0.025%); SCBT, Santa Cruz Biotechnology

**Table 2.**
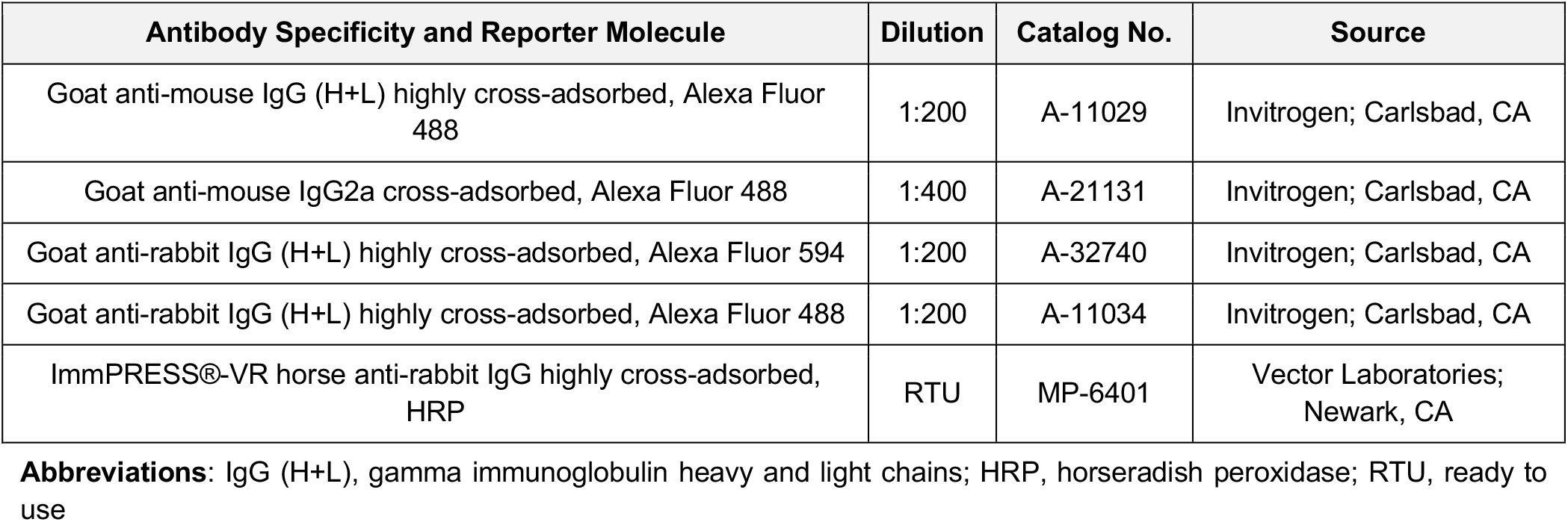
Fluorophore- and Enzyme-Conjugated Secondary Antibodies Used for Immunostaining.

|  |  |  |  |
| --- | --- | --- | --- |
| Antibody Specificity and Reporter Molecule | Dilution | Catalog No. | Source |
| Goat anti-mouse IgG (H+L) highly cross-adsorbed, Alexa Fluor 488 | 1:200 | A-11029 | Invitrogen; Carlsbad, CA |
| Goat anti-mouse IgG2a cross-adsorbed, Alexa Fluor 488 | 1:400 | A-21131 | Invitrogen; Carlsbad, CA |
| Goat anti-rabbit IgG (H+L) highly cross-adsorbed, Alexa Fluor 594 | 1:200 | A-32740 | Invitrogen; Carlsbad, CA |
| Goat anti-rabbit IgG (H+L) highly cross-adsorbed, Alexa Fluor 488 | 1:200 | A-11034 | Invitrogen; Carlsbad, CA |
| ImmPRESS®-VR horse anti-rabbit IgG highly cross-adsorbed, HRP | RTU | MP-6401 | Vector Laboratories; Newark, CA |

**Abbreviations:** IgG (H+L), gamma immunoglobulin heavy and light chains; HRP, horseradish peroxidase; RTU, ready to use

**Table 3.** Fluorophore- and Enzyme-Conjugated Lectins Used for Histochemical Staining.

|  |  |  |  |  |
| --- | --- | --- | --- | --- |
| **Lectins (Acronyms)** | **Carbohydrate Specificity** | **[Lectin]** | **Catalog No.** | **Source** |
| GSL I- isolectin B4 (GSL-IB4), Alexa Fluor 594 | a-D-galactose | 10 µg/ml | I21413 | Invitrogen; Carlsbad, CA |
| GSL I- isolectin B4 (GSL-IB4), Alexa Fluor 488 | a-D-galactose | 20 µg/ml | I21411 | Invitrogen; Carlsbad, CA |
| Lens Culinaris agglutinin (LCA), fluorescein | a-D-mannose, a-D-glucose | 20 µg/ml | L32475 | Invitrogen; Eugene, OR |
| Sambucus Nigra agglutinin (SNA), fluorescein | a-2,6-linked Gal/GalNAc sialic acid residues | 10 mg/ml | L32479 | Invitrogen; Eugene, OR |

**Abbreviations:** α, alpha; Gal/GalNAc, galactose/N-acetyl-galactosamine; GSL I, Griffonia Simplicifolia Lectin I

For immunofluorescence staining, the incubation with primary antibodies was conducted for 1.5 hours at +37°C in a moist chamber followed by labeling with fluorophore-conjugated secondary antibodies for 45 minutes at +37°C in a moist chamber. In the staining where fluorophore-labeled lectins were used, all washing steps were done with PBS containing calcium and magnesium ions. The sections were then cover slipped with ProLong Gold antifade mounting medium (cat. no. P36931; Molecular Probes, Eugene, OR) containing DAPI (4’, 6-Diamidino-2-Phenylindole) to counterstain the nuclei.

For immunohistochemistry, the sections were pretreated with 0.3% hydrogen peroxide solution in distilled water for 15 minutes at room temperature to block endogenous peroxidase activity. The incubation with primary antibodies was conducted for 1.5 hours at +37°C in a moist chamber followed by labeling with horseradish peroxidase (HRP)-conjugated secondary antibodies for 30 minutes at +37°C in a moist chamber. The HRP-conjugated antibodies were visualized using DAB (3,3’-diaminobenzidine) substrate (cat. no. SK-4103; ImmPACT DAB EqV Peroxidase Substrate kit; Vector Labs, Burlingame, CA) at room temperature. The nuclei were counterstained with Vector hematoxylin (cat. no. H-3401; Vector Labs, Burlingame, CA). Omission of primary antibodies served as negative controls.

### 2.3. Microscopy

The stained sections were examined under a Leica DM4000 B microscope (Leica Microsystems, Deerfield, IL) using 20x, 40x and 63x objectives. The images from H&E, Masson’s trichrome, and immunohistochemically stained sections were captured onto a computer with a Leica K3C digital color camera (Leica Microsystems, Deerfield, IL), whereas the double and triple fluorescence labeled images were captured with a Leica K3M digital monochrome camera (Leica Microsystems, Deerfield, IL) using a Leica Application Suite X software. The final figures were digitally assembled from the captured images using Adobe Photoshop CC software (Adobe Systems, San Jose, CA).

## 3. RESULTS

### 3.1. Distribution of CD34+ SC/TCs in non-infarcted myocardium

In remote, non-infarcted myocardium, CD34+ SC/TCs were found in the vessel adventitia, the interstitium between cardiac myocytes (CMs), and alongside the microvessels (Figure 1).

**Figure 1.**
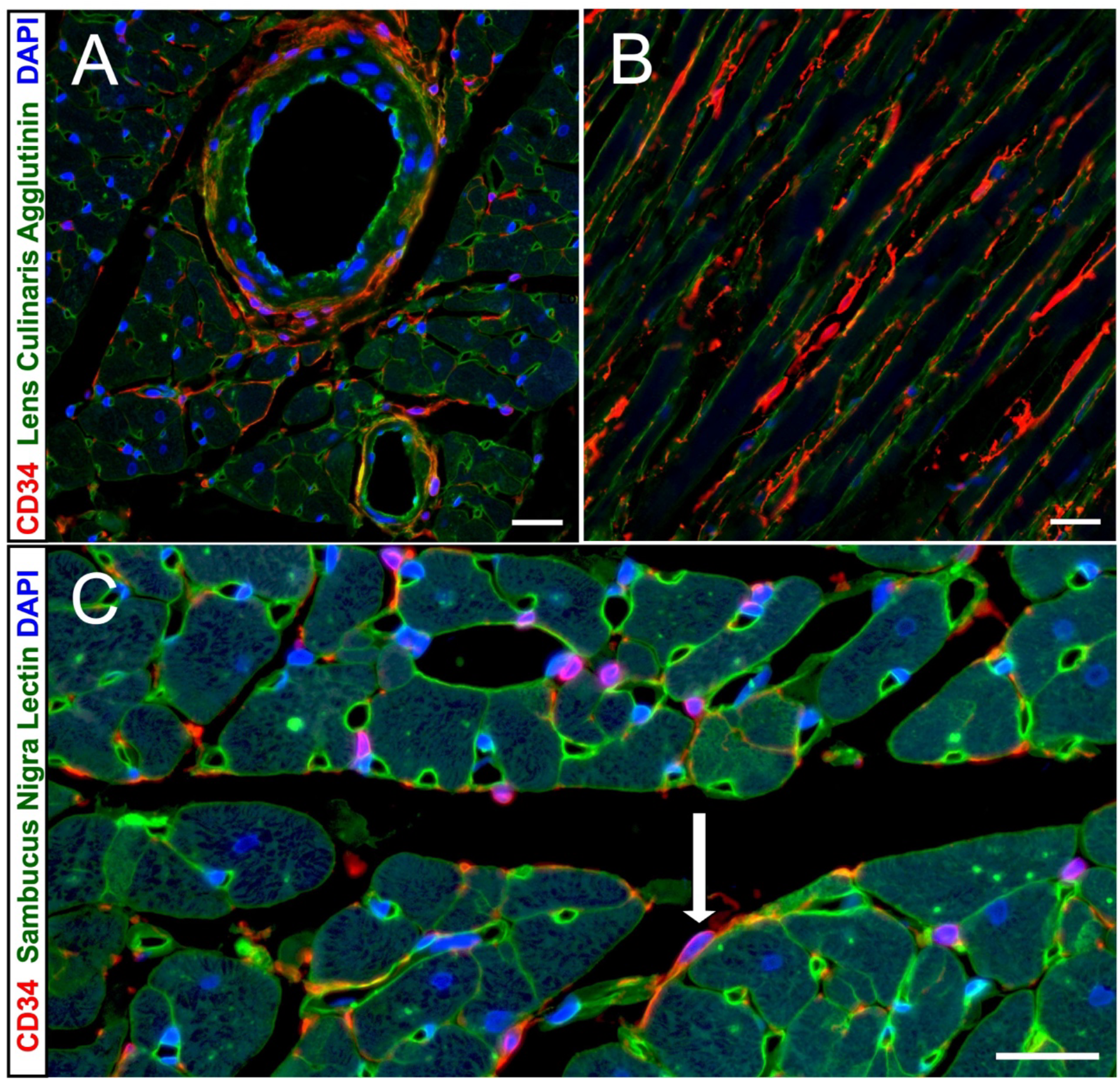
Spatial distribution of CD34+ SC/TCs in remote, non-infarcted myocardium. Representative micrographs from transversely-(**A** and **C**) and longitudinally-(**B**) sectioned areas of remote LV myocardium 3 days after an MI demonstrate the distribution of SC/TCs labeled with an anti-CD34 antibody (red) and the basal lamina of cardiac myocytes and the vessels revealed by Lens Culinaris Agglutinin staining (**A** and **B**) or Sambucus Nigra Lectin (**C**) (green); DAPI is used in all images for a nuclear counterstain (blue). (**A**) CD34+ SC/TCs are present in the vessel adventitia and in the interstitial space between cardiac myocytes. (**B**) CD34+ SC/TCs are seen along the length of cardiac myocytes, often following capillaries. **(C**) This micrograph depicts a resident CD34+ cell within a stroma of intact myocardium that display a small, ovoid cell body with multiple long, thin cytoplasmic processes which are typical phenotypic characteristics of telocytes (white arrow). **Scale bars: 20 µm**

### 3.2. Spatiotemporal Distribution of CD34+ SC/TCs in Developing post-MI Scar

#### 3.2.1. Late Inflammatory Phase of the Healing Process

Three days after an MI, CD34+ SC/TCs were not detectable within the area of coagulative necrosis but were present within the interstitial spaces around surviving cardiac myocytes (CMs) adjacent to the necrotic zone and in the adventitia of residual coronary vessels of the infarcted region (Figure 2).

**Figure 2.**
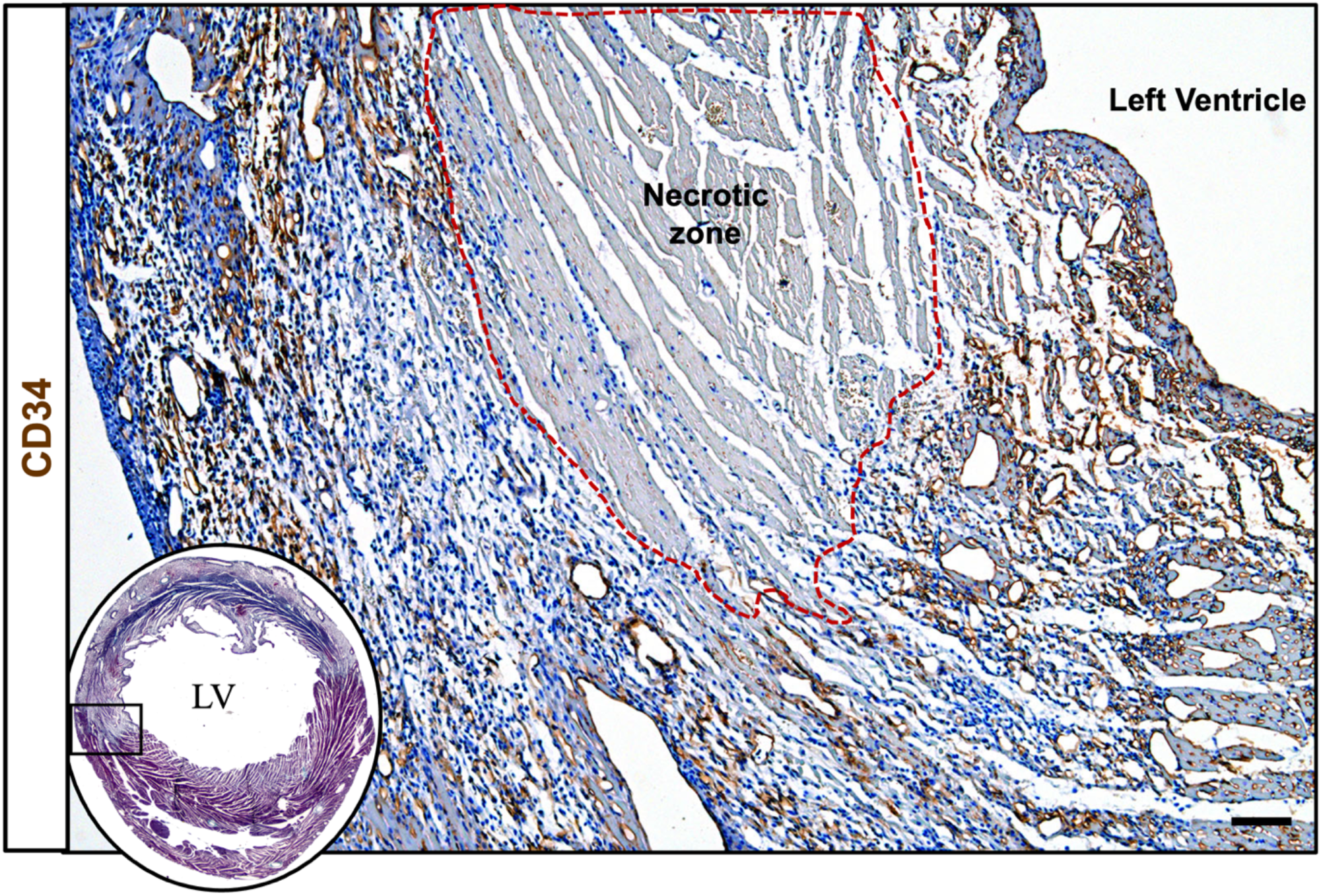
Spatial distribution of CD34+ SC/TCs within the infarcted and peri-infarcted regions during the late inflammatory phase. A representative micrograph demonstrates the tissue distribution of CD34+ SC/TCs visualized with an anti-CD34 antibody that is revealed using HRP-DAB chromogenic reaction (brown). Nuclei are counterstained with Vector hematoxylin (blue). Note that CD34+ SC/TCs are absent in the necrotic zone (outlined by a red-dashed line) but remain visible in the peri-necrotic regions. **Scale bar: 100 µm**

Moreover, CD34+ SC/TCs appeared to be intermingled with phagocytic macrophages involved in the clearing the necrotic debris within the resorption zone, adjacent to the area of coagulative necrosis (Figure 3).

**Figure 3.**
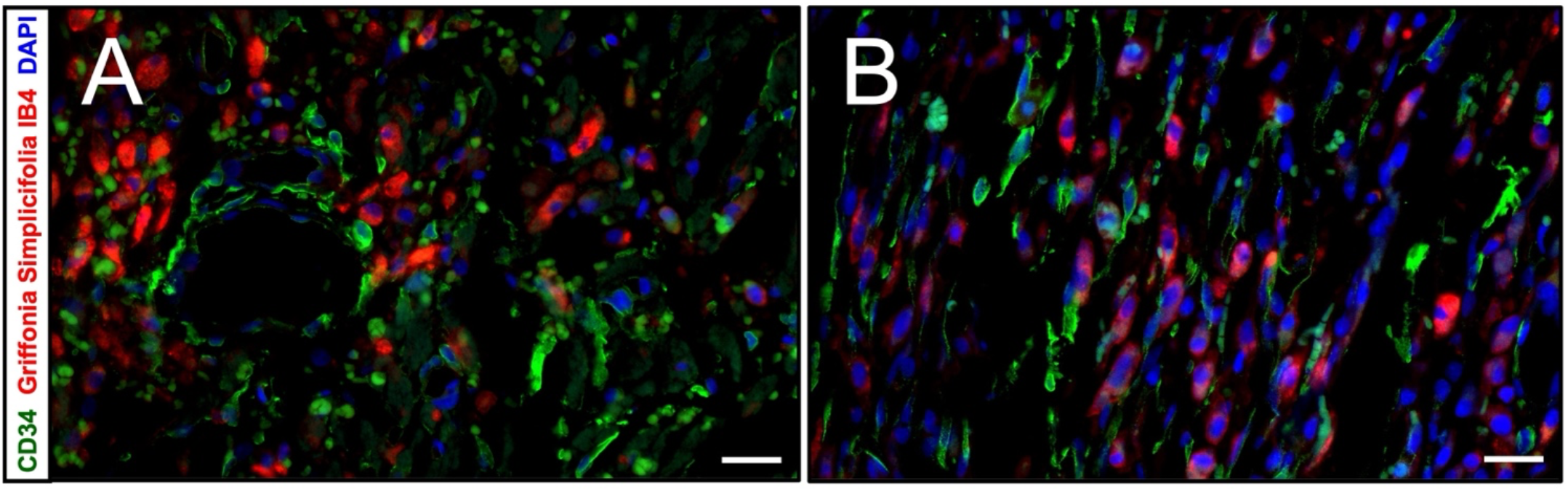
CD34+ SC/TCs and phagocytic macrophages coexist in the areas of necrotic tissue resorption. Representative micrographs display CD34+ SC/TCs (green) intermixed with the phagocytic GS-IB4+ macrophages (red) at the lateral edge of the infarcted region (**A**) and near the endocardium (**B**). DAPI is used for nuclear counterstain (blue). **Scale Bars: 20 µm**

Importantly, the presence of activated macrophages did not deter CD34+ SC/TCs from proliferation as confirmed by the presence of BrdU labeling in the cells (Figure 4A).

**Figure 4.**
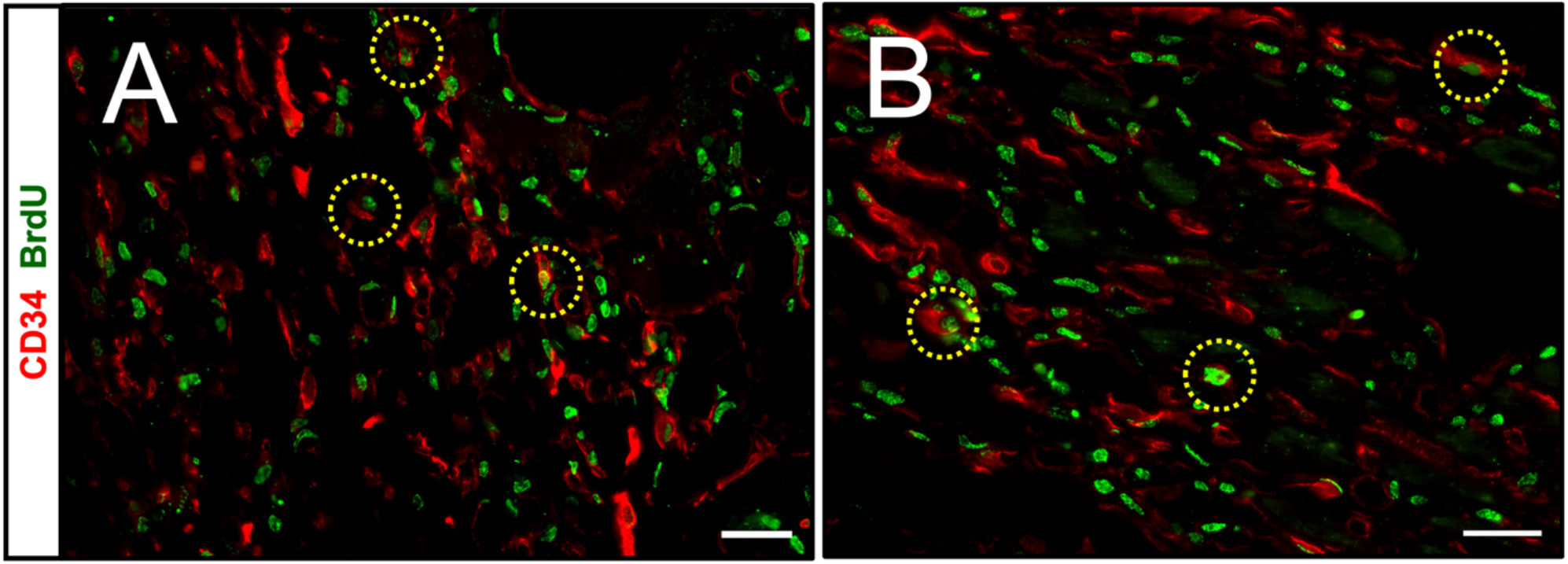
CD34+ SC/TCs present in the developing scar undergo proliferation. Representative images demonstrate the BrdU labeled nuclei (green) in CD34+ SC/TCs (red) in 3-day-old (**A**) and 7-day-old (**B**) MI. Examples of proliferating CD34+ SC/TCs are outlined by the dotted, yellow circles. **Scale Bars: 20 µm**

#### 3.2.2. Early Reparative Phase of the Healing Process

Seven days after MI, a population of CD34+ SC/TCs continued to expand by proliferation and were seen to be spreading throughout the resorption zone (Figure 4B). Surprisingly, the expanded CD34+ SC/TC population seemed to lag behind the macrophages, which were advancing towards the necrotic core, leaving behind the empty basement membranes of resorbed CMs (Figure 5).

**Figure 5.**
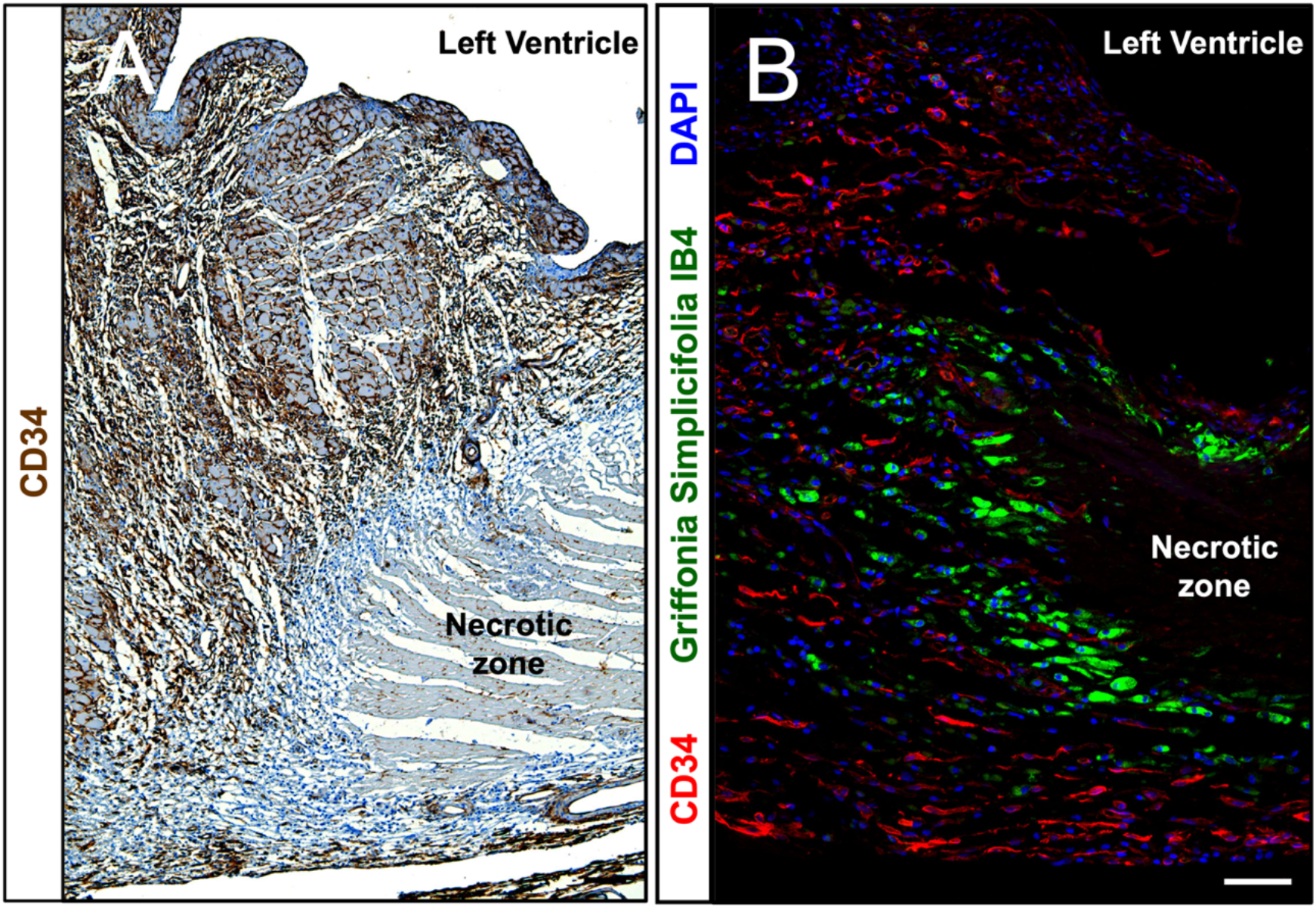
Spatial distribution of CD34+ SC/TCs and phagocytic macrophages in the resorption area during the early reparative phase. These two low-power images were obtained from serial sections of the same peri-infarct region. (**A**) A representative immunohistochemistry micrograph demonstrates the tissue distribution of CD34+ SC/TCs visualized with an anti-CD34 antibody that is revealed by using HRP-DAB chromogenic reaction (brown). Nuclei are counterstained with Vector hematoxylin (blue). Note, the CD34+ SC/TCs remained separated from the necrotic zone by the area occupied only by the hematoxylin-stained nuclei, presumably from macrophages. (**B**) A representative immunofluorescence micrograph shows the gathering of CD34+ SC/TCs (red) that lingered slightly behind the phagocytic GS-IB4+ macrophages (green), which are seen adjacent to the necrotic zone. DAPI is used to counterstain the nuclei. **Scale Bars: 100 µm (A) and 50 µm (B)**

#### 3.2.3. Late Reparative Phase of the Healing Process

Fourteen days after MI, CD34+ SC/TCs were found to be distributed throughout all areas of the developing scar (Figure 6) with exception for a few distinct regions.

**Figure 6.**
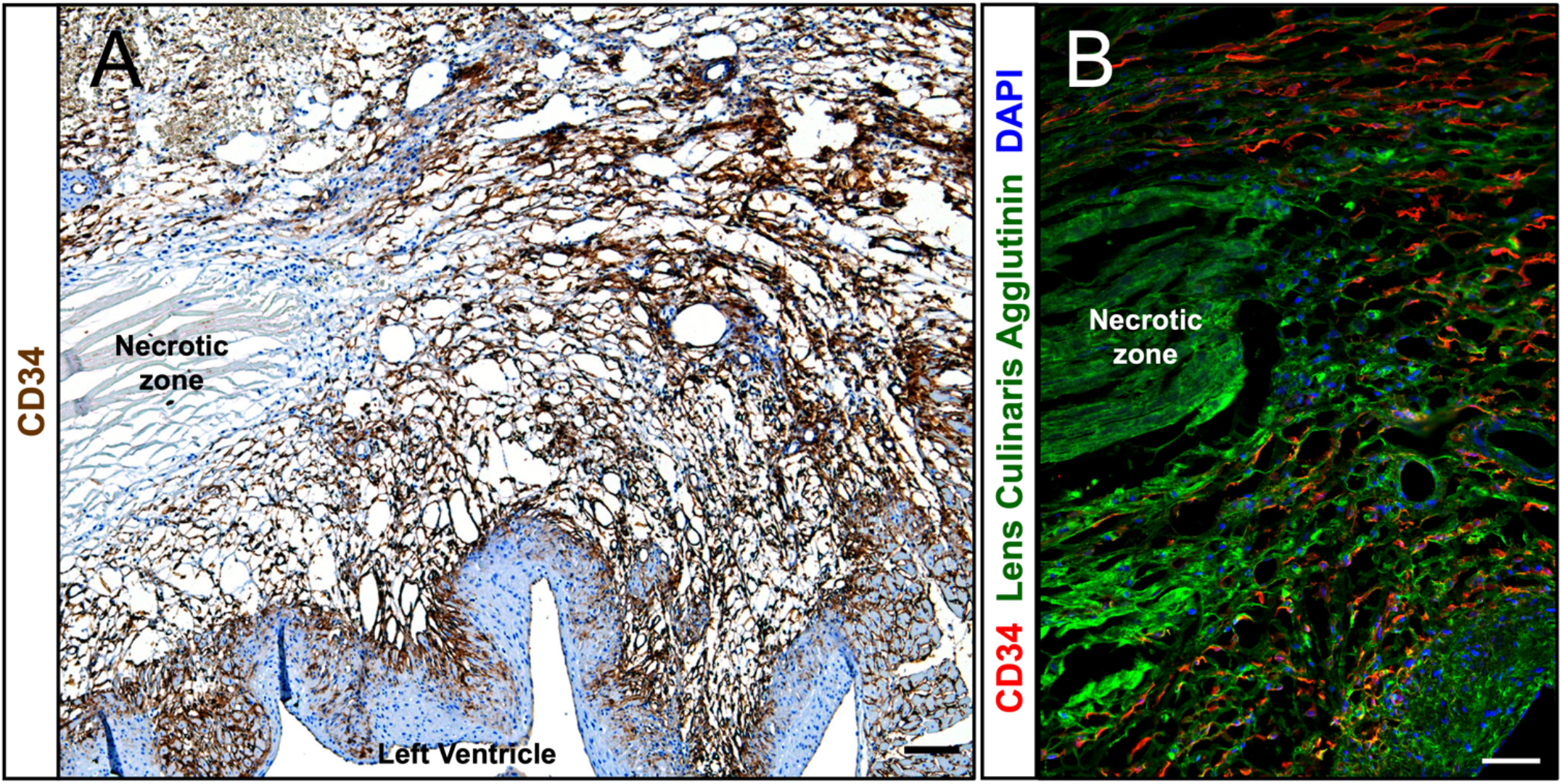
Spatial distribution of CD34+ SC/TCs within the infarcted and peri-infarcted regions during the late reparative phase. These two images were obtained from serial sections of the same peri-infarct region. (**A**) A representative immunohistochemistry micrograph demonstrates the tissue distribution of CD34+ SC/TCs labeled with an anti-CD34 antibody that is revealed by using HRP-DAB chromogenic reaction (brown). Nuclei are counterstained with Vector hematoxylin (blue). Note that CD34+ SC/TCs distributed throughout the entire area of resorption, reaching the necrotic core. (**B**) A representative immunofluorescence micrograph shows that CD34+ SC/TCs (red) were predominantly found in the spaces between basement membranes of resorbed cardiac myocytes, revealed by Lens Culinaris Agglutinin (LCA) staining (green). **Scale Bars: 100 µm**

Interestingly, in addition to the residual necrotic core (Figure 6), CD34+ SC/TCs were absent from areas occupied by cells expressing alpha smooth muscle actin (myofibroblasts and smooth muscle cells), such as granulation tissue, the tunica media of residual coronary arteries, and the subendocardial fibroelastic thickening (Figure 7).

**Figure 7.**
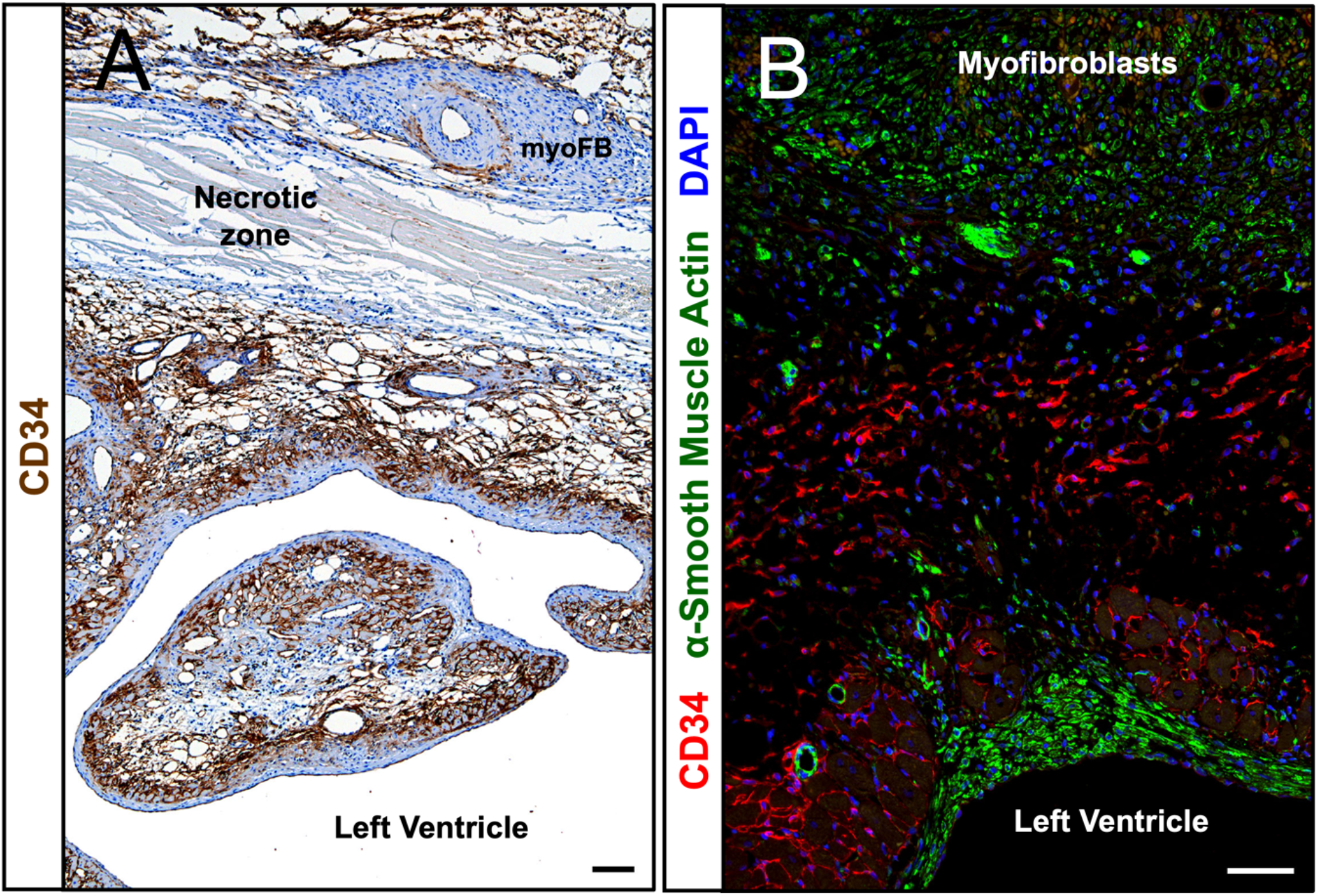
Spatial distribution of CD34+ SC/TCs in regions dominated by alpha smooth muscle actin (SMA) expressing cells. (**A**) SC/TCs stained with an anti-CD34 antibody that is visualized with HRP-DAB chromogen reaction (brown). Nuclei are counterstained with Gill hematoxylin (blue). (**B**) A double immunofluorescence staining with the antibodies againsCD34 (red) and alpha SMA (green) DAPI is used for nuclei counterstain (blue). In both micrographs, CD34+ SC/TCs are absent in areas where alpha SMA-positive cells are present. **Scale Bars: 100 µm (A) and 50 µm (B)**

Furthermore, a dual staining with anti-CD34 antibody and LCA showed that the bodies and long, slender cytoplasmic processes of CD34+ SC/TCs appeared to intimately align with acellular basement membranes, suggesting that these extracellular structures were used as potential scaffolds for CD34+ SC/TC migration (Figure 8).

**Figure 8.**
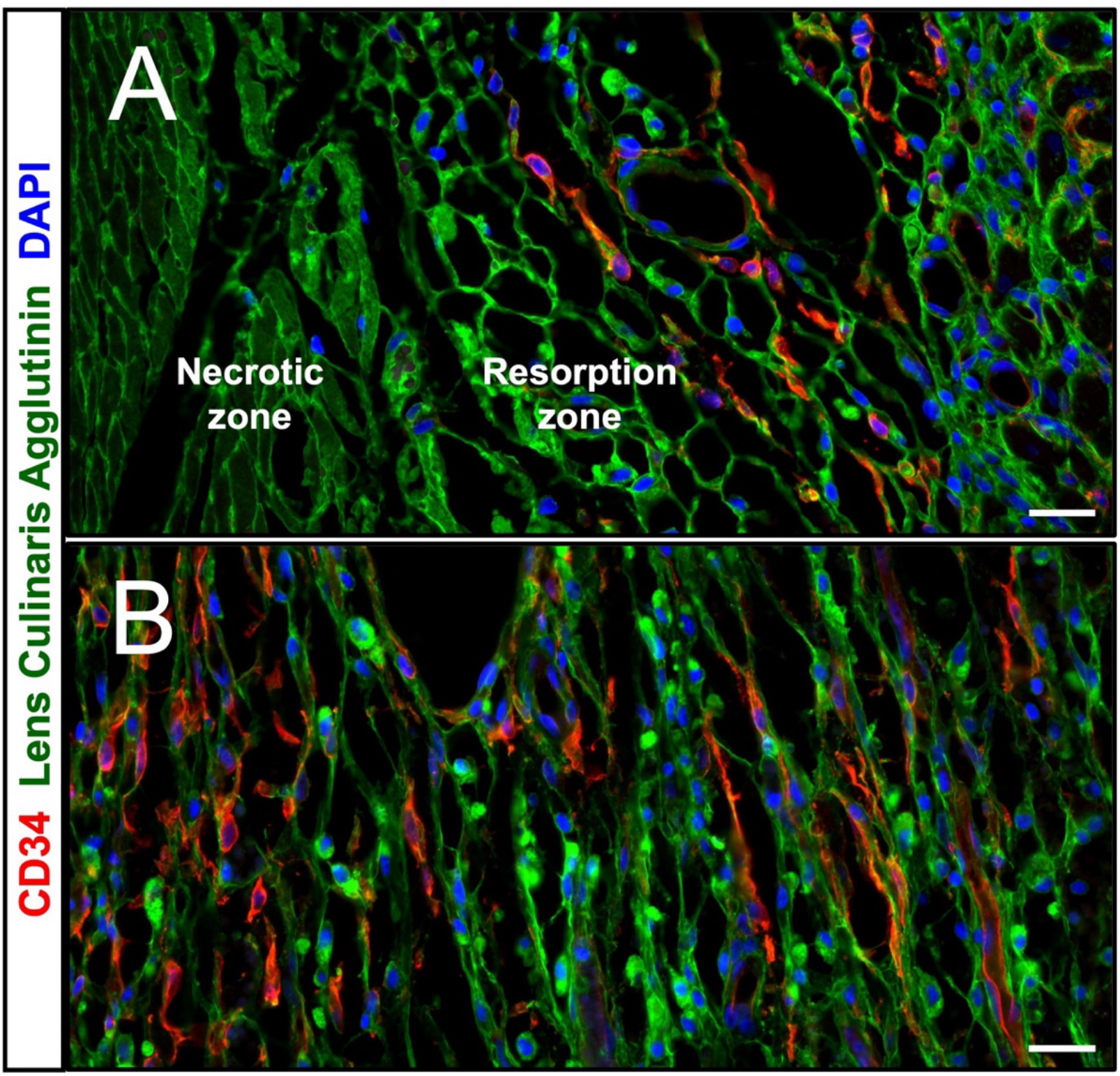
The arrangement of CD34+ SC/TCs along the acellular basement membranes of resorbed cardiac myocytes within the resorption zone. Transverse (**A**) and longitudinal (**B**) views from the resorption zone demonstrate the distribution of CD34+ SC/TCs (red) along the LCA stained basal membranes (green), outlining the empty spaces. Note that in both micrographs the bodies and cytoplasmic extensions of CD34+ SC/TCs appear well approximated to acellular basement membranes of resorbed CMs. **Scale Bars: 20 µm**

In addition, aggregates of flattened CD34+ SC/TCs cells were found at the edges of the scar where they covered the boundary formed by peri-infarcted CMs (Figure 9A) and newly formed capillaries (Figure 9B).

**Figure 9.**
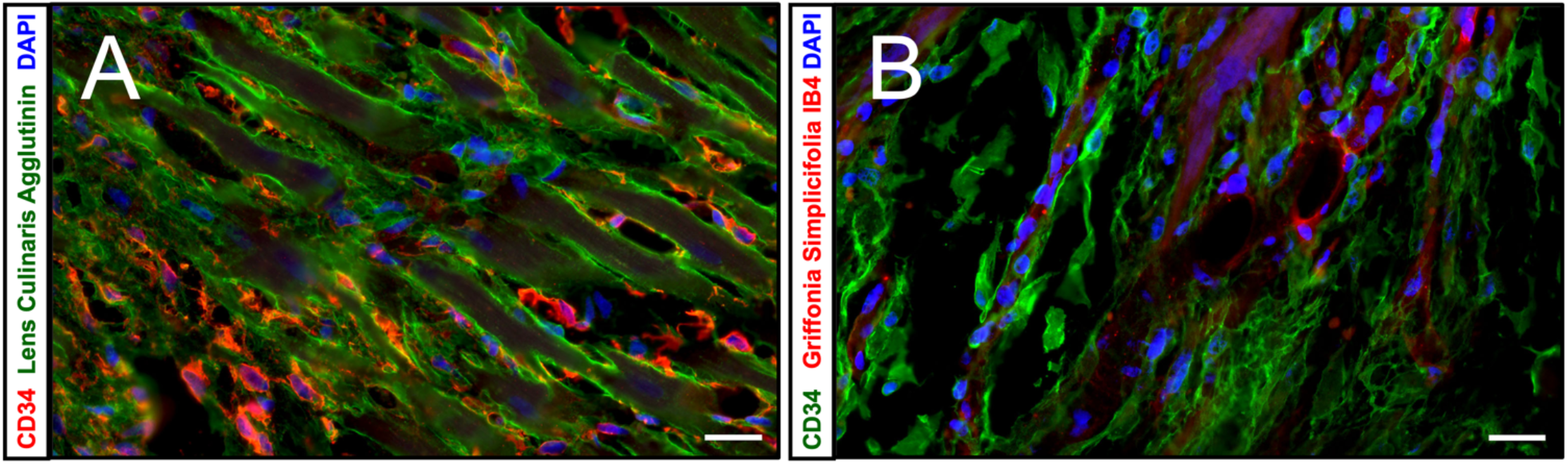
A spatial pattern of CD34+ SC/TCs at the scar margin in 14-day-old MI. (**A**) Flattened and enlarged CD34+ SC/TCs (red) appear to enfold peri-infarcted cardiac myocytes, visualized by staining with Lens Culinaris Agglutinin (green). (**B**) CD34+ SC/TCs (green) are wrapping up the capillaries labeled with GS-IB4 (red) at the age of the MI. In both images, nuclei are counterstained with DAPI (blue). **Scale Bars: 20 µm**

## 4. DISCUSSION

This study, for the first time, demonstrates a population of resident CD34+ SC/TCs following a reproducible, dynamic pattern of distribution within the ischemic region of the heart during the late inflammatory and proliferative phases of post-MI healing. The key findings were as follows: 1) in late inflammatory phase (three days after MI), CD34+ SC/TCs were absent in the necrotic zone but were found intermingled with the phagocytic macrophages in the area of myocardial resorption; 2) in the early proliferative phase (seven days after MI), a population of CD34+ SC/TCs underwent an expansion via proliferation, but appeared to fall behind advancing macrophages which were clearing the cellular debris and preserving the basal membranes of resorbed cardiac myocytes; and 3) in the late reparative phase (fourteen days after MI), CD34+ SC/TCs were found to spread throughout the infarcted region, presumably by migrating along acellular basement membranes toward the necrotic core. They were further observed to engulf the surface of surviving cardiac myocytes and newly formed microvessels at the scar margin. Most interesting, CD34+ SC/TCs were absent from regions occupied by cells expressing alpha SMA, notably myofibroblast-rich granulation tissue, the media of the residual arteries, and the subendocardial fibroelastic thickening.

### 4.1. Myocardial CD34+ SC/TCs Serve as a Modulators for Scar Tissue Organization and Reparative Angiogenesis

This study has confirmed the association of CD34+ SC/TCs with the stromal components of both intact and post-infarcted myocardial tissue. According to previously published reports, these cells could be involved in several reparative and fibrotic mechanisms across various organ systems and disease spectrums. Proposed mechanisms have included the ability of CD34+ SC/TCs to serve either as an activator of various stem cells^15^ or as progenitor cells assisting with mesenchymal tissue repair^16^. Furthermore, some authors have concluded that these cells could function as organizers of the tissue architecture^17^ while others have argued that the intricate network they form within the tissue interstitial space is serving as a paracrine-based communication network^18-22^.

Furthermore, in accord with earlier reported observations suggesting that CD34+ SC/TCs could be the source of miRNA exosomes promoting angiogenesis^21^, it was found that in post-MI myocardium these cells were often in close proximity to newly-formed capillaries which extend from the peri-infarcted border into the healing region, implying their potential role in paracrine signaling. However, these results do not exclude the possibility that CD34-expressing cells observed around new microvessels are not bone-marrow derived endothelial cell precursors, which are needed to form capillaries in healing tissue^23^.

The current understanding of CD34+ SC/TCs function is wildly varied, and while all these mechanisms must be considered, the evidence presented in this study corroborates that these cells are involved in extremely close interactions with interstitial components of intact and healing myocardium.

### 4.2. Myocardial CD34+ SC/TCs and Phagocytic Macrophages Appeared to Closely Collaborate During Tissue Debris Clearing and Cell Migration

Macrophages are well known to be involved in repair and regeneration of damaged tissues within the body. However, they are not only responsible for clearing of necrotic debris but have been also shown to play an important role in extracellular matrix turnover in both healthy and disease conditions. In general, macrophages can be subdivided into two types: M1 and M2. M1 macrophages are primarily responsible for inducing inflammation and promoting clearance of damaged tissue while M2 macrophages are mainly recognized as the anti-inflammatory cells which promote tissue regeneration and angiogenesis. Tissue repair and regeneration following an MI involves both subtypes of macrophages. The M1 macrophages have been demonstrated to peak in quantity around three days following MI, whereas M2 macrophages peak around seven days post-MI, with both types diminishing thereafter^24^.

The relationship between CD34+ SC/TCs and macrophages is not well understood, and there is clear variation in their function depending on the context. For example, in periodontitis, telocytes have been implicated in the transition of M1 macrophages to hybrid M1/M2 macrophages through hepatocyte growth factor (HGF) signaling pathways^26^. Conversely, in a mouse model, telocytes were shown to induce M1 differentiation through NF-KB pathways in cases of endometriosis^27^. CD34+ SC/TC and macrophage interactions have not been well described in the heart.

This study reveals that in the late inflammatory phase following MI, proliferating CD34+ SC/TCs were seen intermingled with prophagocytic, presumably type M1, macrophages, suggesting the ability of these two types of cells to coexist within the proinflammatory milieu. Furthermore, in the reparative phase, CD34+ SC/TCs appeared to trail slightly behind the advancing macrophages and using the intact basement membranes left from the resorbed cardiac myocytes as the routes for their migration throughout all regions cleared from tissue debris.

### 4.3. CD34+ SC/TCs and Alpha-SMA+ cells, Presumably Myofibroblasts, Seemed to Favor the Different Mechanisms of Tissue Healing

This study demonstrated a clear distinction in spatial distribution of CD34+ SC/TCs and the cells expressing alpha-SMA, namely myofibroblasts and vascular smooth muscle cells, within healing myocardium since neither cell types occupied the same spaces as the other. Most interesting, these findings corroborated similar observations in multiple other organ systems and diseases^28-30^.

Prior studies have already investigated the relationship between these cell types during tissue repair and fibrosis. For example, some have suggested that the phenotypic conversion of fibroblasts to myofibroblasts withing growing granulation tissue could be partially mediated by CD34+ SC/TCs^31,32^. Alternatively, some authors have proposed that CD34+ SC/TCs could directly undergo phenotypic conversion to myofibroblasts^33^. However, this mechanism seems less likely, given that none of the CD34+ SC/TCs in this study co-expressed alpha-SMA when dual staining with CD34, indicating a lack of cells transitioning between phenotypes. Considering the abovementioned findings that CD34+ SC/TCs and alpha-SMA+ cells almost never occupied the same areas within healing tissue, it is reasonable to propose that the milieu existing in the areas occupied by alpha-SMA+ cells could repel CD34+ SC/TCs from these regions. Although our current data cannot provide a conclusive explanation for this phenomenon, they confirm a unique relationship between these cell types is present during the development of granulation tissue and scar formation.

### 4.4. Conclusion and Future Direction

Altogether, the above findings support the conclusion that CD34+ SC/TCs play a vital role in post-MI healing. Accordingly, these cells should be considered in future studies as a potential target for scar modifying therapies, with special attention paid to their interactions and associations with cell types implicated early in the repair process.

### 4.5. Study Limitations

The current investigation into the role of CD34+ SC/TCs during post-MI repair process remained somewhat deficient because there is no consensus on how to properly identify these cells. Some investigators suggested the exclusive utilization of electron microscopy to confirm the unique morphology of these cells on high resolution images, when others suggested the use of immunophenotypic characterization by exploiting combinations of various immunoprobes, such as antibodies against CD34, PDGFR alpha, CD117 and many others. However, all these methodological approaches have their own limitations. For example, the use of electron microscopy markedly reduces the size of tissue samples, which may interfere with assessment of overall spatial distribution. On the other hand, precise immunophenotypic characterization of these cells is also problematic since it appears to vary across different organ systems or species. In this regard, anti-CD34 antibodies were the most reliable and only consistent marker for cells that represented the morphology of telocytes so far. Almost all previously published reports that we used to conduct our current study have consistently employed an anti-CD34 immunostaining either in combination with other markers or imaging techniques. Accordingly, this study to identified CD34+ SC/TCs utilizing labeling with an anti-CD34 antibody marker in combination with phenotypic cell characterization. It is important to emphasize that since the findings of this study remain largely descriptive, some level of caution must be taken when the true nature of the myocardial CD34+ cells is considered.

## 5. ACKNOWLEDGEMENTS

None.

## 6. FUNDING

This research has been funded in part by a grant from Camden Health Research Initiative. The author declared no potential conflicts of interest with respect to the research, authorship, and/or publication of this article.

## Notes

### Competing Interest Statement

The authors have declared no competing interest.

